# Maternal obesity causes fetal cardiac hypertrophy and alters adult offspring myocardial metabolism in mice

**DOI:** 10.1101/2021.09.15.460457

**Authors:** Owen R. Vaughan, Fredrick J. Rosario, Jeannie Chan, Laura A. Cox, Veronique Ferchaud-Roucher, Karin A. Zemski-Berry, Jane E.B. Reusch, Amy C. Keller, Theresa L. Powell, Thomas Jansson

## Abstract

Obesity in pregnant women causes fetal cardiac dysfunction and increases offspring cardiovascular disease risk but its effect on myocardial metabolism is unknown. We hypothesised that maternal obesity alters fetal cardiac expression of metabolism-related genes and shifts offspring myocardial substrate preference from glucose towards lipids. Female mice were fed control or obesogenic diets before and during pregnancy. Fetal hearts were studied in late gestation (embryonic day, E18.5; term≈E21) and offspring were studied at 3, 6, 9 or 24 months postnatally. Maternal obesity increased heart weight and peroxisome proliferator activated receptor γ (*Pparg*) expression in female and male fetuses and caused left ventricular diastolic dysfunction in the adult offspring. Cardiac dysfunction progressively worsened with age in female, not male, offspring of obese dams, compared to age-matched controls. In 6-month-old offspring, exposure to maternal obesity increased cardiac palmitoyl carnitine-supported mitochondrial respiration in males and reduced myocardial ^18^F-fluorodeoxyglucose uptake in females. Cardiac *Pparg* expression remained higher in adult offspring of obese than control dams and correlated with contractile and metabolic function. Maternal obesity did not affect cardiac palmitoyl carnitine respiration in females or ^18^F-fluorodeoxyglucose uptake in males, or alter cardiac ^3^H-oleic acid uptake, pyruvate respiration, lipid content or fatty acid/glucose transporter abundance in offspring of either sex. The results support our hypothesis and show that maternal obesity affects offspring cardiac metabolism in a sex-dependent manner. Persistent upregulation of *Pparg* expression in response to overnutrition *in utero* may mechanistically underpin programmed cardiac impairments and contribute to cardiovascular disease risk in children of women with obesity.

## INTRODUCTION

The global prevalence of obesity in women of reproductive age is rapidly increasing ^1^. Obesity in pregnant women predisposes their children to metabolic syndrome and a range of non-communicable diseases throughout life, thereby placing a substantial burden on the health of the population ^2^. Children of women who have obesity during pregnancy are at 30% greater risk for cardiovascular disease in adulthood ^3^. Diet-induced obesity in pregnant animals similarly leads to abnormal cardiovascular function in their adult offspring, irrespective of postnatal diet ^4, 5^, demonstrating that lifelong phenotype is programmed before birth. However, the mechanisms underlying cardiometabolic programming by maternal obesity remain poorly understood and neither the specific programming signals nor their primary targets in the fetus have been clearly identified.

Fetuses of obese women have cardiac hypertrophy and contractile dysfunction as early as the first trimester ^6, 7^, and these abnormalities persist at least into childhood ^8^. Similarly, in experimental animals, maternal obesity increases fetal heart size, wall thickness, cardiomyocyte size, inflammation and collagen content, and impairs contractility ^9–12^. These data suggest that maternal obesity directly affects the heart *in utero*.

Maternal obesity alters placental transport and fetal delivery of glucose and lipids, in both humans and experimental animals ^13–16^. Maternal obesity also increases fetal cardiac lipid storage in sheep ^11^, and impairs glucose uptake and mitochondrial respiration in isolated cardiomyocytes from the offspring in rodents ^17, 18^. Impaired diastolic function in people with diabetes is associated with increased cardiac fatty acid uptake and oxidation, but reduced glucose uptake ^19, 20^ whilst genetic modifications that increase cardiac lipid uptake in mice cause contractile dysfunction ^21^. Whether abnormal cardiac glucose and lipid metabolism contributes to cardiac dysfunction in the offspring of pregnancies complicated by maternal obesity remains unknown.

We have developed a mouse model of maternal obesity associated with fetal overgrowth that closely replicates the phenotype of obese pregnant women in terms of maternal physiology, placental nutrient transport and fetal growth ^13^. In this model, administration of adiponectin to increase maternal concentrations of this metabolic hormone in obese dams, to the levels observed in control animals, normalises placental nutrient transport, prevents fetal overgrowth and mitigates cardiac diastolic dysfunction in the adult offspring ^22, 23^. In the current study, we used our established model to determine the effect of maternal obesity on the fetal cardiac transcriptome and on cardiac contractile and metabolic function in the adult offspring. We hypothesised that maternal obesity alters fetal cardiac expression of metabolism-regulating genes and increases lipid metabolism but reduces glucose metabolism in hearts of adult offspring. We also measured cardiac histone acetylation, which is linked to heart failure ^24^ and is sensitive to nutrient availability ^25^, to determine whether it plays a role in fetal cardiac programming by maternal obesity.

## MATERIALS AND METHODS

### Animals

All procedures were conducted with approval from the Institutional Animal Care and Use Committee of the University of Colorado. Female C57BL6/J mice, proven breeders, were fed *ad libitum* with an obesogenic diet (Ob, n=31) consisting of high fat pellets (Western Diet D12079B, 41 kcal% fat) supplemented with sucrose solution (20 %), or a control diet (Con, D12489B, 10.6 kcal% fat, n=50). All animals had *ad libitum* access to fresh water and were housed under standard 12hr: 12hr dark:light conditions. When females fed the Ob diet had gained 25% of their initial body weight, they were mated overnight with stud males. Age-matched Con females were mated simultaneously. Successful mating was confirmed by the presence of a copulatory plug the following morning, designated embryonic day (E) 0.5 (term ∼E19.5). Pregnant females were subsequently housed in pairs. On E18.5, a subset of Con (n=5) and Ob (n=5) dams were euthanised by CO_2_ asphyxiation and cervical dislocation. The uterus was exposed, fetuses were excised and weighed then their hearts were dissected, weighed and snap frozen in liquid N2. A fetal tail snip was also collected for subsequent determination of sex, by *Zfy-1* genotyping ^26^. The remaining Con and Ob dams delivered naturally at term and continued to consume their respective diets throughout lactation, until all pups were weaned onto standard chow at age 4 weeks. Pups were subsequently housed in same sex groups, from multiple litters. In total, 187 offspring were used in the study.

### Experimental procedures

#### Echocardiography

Cardiac structure and function were assessed using transthoracic echocardiography in a subset of offspring at both 3 and 6 months of age (females n=10 Con, 6 Ob; males n= 7 Con, 7 Ob), as described in Supplemental Methods. One week after the second echocardiography assessment, offspring were fasted for 4 hr then euthanised by CO_2_ asphyxiation and cervical dislocation. Hearts were perfused with PBS then excised and weighed. The left ventricle was dissected and snap frozen in liquid N2 for gene expression analyses. Thirty-nine additional animals that had not undergone echocardiography were also euthanised at age 6 months, and tissues collected in the same manner for gene expression and lipidomic analyses.

Separate cohorts of offspring underwent echocardiographic assessment by the same operator at ages 9 months (n=28) or 24 months (n=22), without being imaged at earlier time points. These animals were euthanised seven days after echocardiography and their tissues collected for use in other studies.

#### Positron emission tomography

Cardiac glucose uptake was assessed *in vivo* in forty-eight 6-month old offspring using positron emission tomography (PET) with ^18^F-fluorodeoxyglucose (^18^F-FDG) tracer. Animals were fasted for 4 hours prior to the study then anaesthetised (2% isoflurane, inhaled) and positioned in dorsal recumbency on a warming pad at 37°C. Animals were then placed inside a PET imaging system (Inveon microPET, Siemens Medical, Knoxville, TN, USA) and a bolus dose of ^18^F-FDG was administered intravenously, to the tail vein (250uCi). Fifteen dynamic PET image frames were collected at successive timepoints over a period of 35 minutes. A whole-body computerised tomography image, without contrast agent, was subsequently collected to confirm anatomical distribution of the ^18^F-FDG tracer. Mice were removed from the scanner, recovered from anaesthesia and housed in a lead-shielded cage with *ad libitum* access to food and water, before being returned to their home cage 24 hr later.

Ten animals that underwent PET imaging were excluded from the subsequent analysis because (a) they did not tolerate anaesthesia, (b) poor tail vein patency hindered tracer infusion, or (c) movement interfered with collection of dynamic frame images. Dynamic frame PET images were assembled for each animal and voxel intensities (radioactivity) determined for manually defined regions of interest in the left ventricular myocardium (study compartment) and lumen (reference compartment). Radioactivity at each timepoint, within each compartment, was corrected for decay, animal weight and the amount of tracer injected. The ratio of radioactivity in the left ventricular myocardium to that in the reference compartment was then plotted against normalised time from tracer injection, according to the Patlak method ^27^. Separate linear regression lines were fitted to data from Con and Ob groups and the slopes compared by extra sum-of-squares F test, within each sex. The slope of the Patlak plot represents the clearance of ^18^F-FDG from the blood, into the myocardium.

This group of animals was euthanised seven days after imaging, as described above. A portion of the left ventricle was placed into ice-cold biopsy preservation solution (BIOPS ^28^) for analysis of mitochondrial respiration within ∼3 hours.

#### *In vivo* left ventricular ^3^H-oleic acid uptake

Cardiac fatty acid uptake was quantified in 20 six-month old offspring using a radioactive tracer, as described previously ^29^. Briefly, 4.5µCi ^3^H-oleic acid (NET289001MC, Perkin Elmer), was dried down under N2 then resuspended in PBS and combined 1:1 with 40% fatty-acid free bovine serum albumin to a final volume of 200µl, at 37°C with shaking. Mice were anaesthetised with ketamine and xylazine (*i.p.*), the tail vein canulated and the BSA-complexed ^3^H-oleic acid tracer delivered as a bolus. Up to five minutes later, mice were euthanised with sodium pentobarbital solution (*i.v.*), rapidly exsanguinated and the heart perfused with PBS. Plasma was separated from blood by centrifugation. Left ventricles (LV) were dissected, weighed then digested for 48 hr at 50°C in Biosol (National Diagnostics). LV and plasma radioactivity were determined by liquid scintillation counting and ^3^H-oleic acid clearance calculated from the ratio of total LV counts to the area under the curve of plasma counts versus time from tracer infusion, divided by LV weight.

### Biochemical and molecular analyses

#### RNA extraction, sequencing and Ingenuity Pathway Analysis

Heart tissues were collected from male and female fetuses within litters from Con (n=5) and Ob (n=5) dams and stored at -80°C. Individual frozen tissue samples (∼10 mg) were homogenized in TRI Reagent and total RNA was isolated using Direct-zol RNA MiniPrep kits (Zymo Research, Irvine, CA) according to the manufacturer’s instructions. After eluting RNA from the Zymo-Spin column in 50 µl DNase/RNase-free water, RNA quality was assessed using an Agilent 2100 Bioanalyzer (Agilent Technologies, Inc., Santa Clara, CA). RNA concentration was quantified using Qubit RNA HS assay kits and a Qubit 2.0 Fluorometer (Thermo Fisher Scientific, Wilmington, DE). Total RNA was stored at -80°C. Differentially expressed genes between Con and Ob fetuses were identified and functionally annotated, as described in the Supplementary Methods.

#### High resolution *in situ* respirometry

Carbohydrate- and lipid supported rates of mitochondrial respiration were assessed in cardiac muscle from 6-month old offspring, using high resolution respirometry. Biopsies of fresh left ventricular tissue in ice-cold BIOPS solution were dissected under magnification into myofiber bundles of approximately 1 mg. The sarcolemmal membrane was permeabilized by incubating in saponin solution (40µg ml^-1^, 20 min) then washed, accurately weighed and placed in an oxygraph chamber (Oroboros O2k Respirometer), in respiration medium (MIRO5, ^28^) with blebbistatin (5mM) to inhibit muscle contraction. Medium was equilibrated with O_2_ gas to an initial dissolved concentration of 400µM and the chamber was sealed. Catalase was also added to the respiration medium to allow liberation of oxygen by H_2_O_2_ addition during the experiment, thus retaining dissolved O_2_ concentration between 300 µM and 400 µM.

O_z_ consumption rate was determined during the sequential addition of one of two different combinations of substrates and inhibitors, designed to determine the basal and maximal rates of carbohydrate- and lipid-supported respiration, respectively. In the first assay, pyruvate (5mM) and malate (2mM) were added to the oxygraph chamber to supply the electron transport chain via reduced intermediates generated in the citric acid cycle. In the second assay, palmitoyl carnitine (5µM) and malate (1mM) supplied electrons via mitochondrial β-oxidation. In both cases, leak state oxygen consumption was first measured in the absence of ADP. Then ADP (2mM) was introduced to measure basal respiration rate, coupled to ATP synthase activity. Glutamate (3mM) and succinate (6mM) were subsequently provided to support direct electron flux to complex I and II of the electron transport chain, allowing maximum oxidative phosphorylation capacity to be measured. ATP synthase activity was abolished using oligomycin (4µg/ml), to measure leak state oxygen consumption supported by both fatty acid oxidation and direct complex I/II electron entry. Finally, FCCP (carbonyl cyanide 4-(trifluoromethoxy)phenylhydrazone) was added in increments of 0.5µM to permeabilise the inner mitochondrial membrane, until maximal, uncoupled oxygen consumption was achieved, measuring electron transport chain capacity. Respiration rates were corrected to fresh tissue mass. Coupling efficiencies for each of the two combinations of substrates was calculated as 1 minus the ratio (leak state respiration/maximal electron transport chain capacity).

#### Targeted lipidomic analyses

Cardiac triglyceride contents were determined by colorimetric assay (MAK266, Sigma-Aldrich). Myocardial content of individual fatty acids, ceramides and diacylglycerol species was measured in adult offspring using targeted mass spectrometry, described in Supplementary Methods.

### Western blot

Fatty acid and glucose transporter protein abundance was measured in cardiac tissue from 6-month old offspring using western blotting. Frozen heart samples were homogenised in hepes-tris buffered saline with protease and phosphatase inhibitors (Sigma Aldrich). Samples were then loaded in Laemmli buffer, resolved by polyacrylamide gel electrophoresis and transferred to polyvinylidene fluoride membrane. Membranes were incubated with primary antibodies to CD36, FATP1, FATP6, GLUT1 and GLUT4 then with a horseradish peroxidase linked secondary antibody and visualised using enhanced chemiluminescence reaction and a gel imaging system. Protein abundance was determined by densitometry of specific bands, corrected for protein loading (amido black stain).

### Gene expression

Cardiac expression of selected mRNAs related to lipid metabolism was determined in a targeted manner using qRT-PCR. RNA was extracted from frozen ventricular tissue from 6-month-old Con (females n=9, males n=11) and Ob (females n=10, males n=10) offspring using a commercially available kit (RNeasy Plus mini Kit, Qiagen) then reverse transcribed to cDNA (High Capacity cDNA Reverse Transcription kit, Applied Biosystems). Relative expression of target mRNAs was determined by qRT-PCR using SYBR Green chemistry and forward and reverse primers, as detailed in Supplemental Table 1. The efficiency of all primer pairs was confirmed to be between 80 and 110% by calculating the linear gradient of the relationship between average Ct value and dilution factor for a 5-fold serially diluted standard curve of pooled cDNA. PCR product size was validated by agarose gel electrophoresis. Gene expression was determined relative to the geometric mean of *Rna18s* and *Rps29* expression using the ddCt method.

### Statistics

Results are presented as mean ± SEM. All statistical analyses were conducted separately in female and male offspring. Normality of data was assessed by Shapiro-Wilk test. Echocardiographic measurements of cardiac function and morphometry were analysed by two-way ANOVA, with maternal obesity and postnatal age as independent factors. When there was a significant interaction between these two factors, the simple effect of maternal obesity at each age was assessed by Sidak post-hoc test. For all other measurements, Con and Ob groups were compared by Student’s t-test or by Mann-Whitney test, if data did not conform to a normal distribution. Analyses of lipidomic data were corrected for multiple comparisons using the Holm-Sidak method. Linear relationships between variables were determined by Pearson’s correlation. In all cases, significance was taken at the level P<0.05.

## RESULTS

### Maternal obesity in pregnant mice induces fetal cardiac hypertrophy and transcriptional activation of lipid metabolism genes

Maternal obesity increased fetal heart weight at E18.5, as a percentage of total body weight, in both male and female fetuses (Fig. 1A, B). Absolute heart and body weights were also greater in fetuses of obese dams, compared to controls (Table 1).

**Fig. 1.**
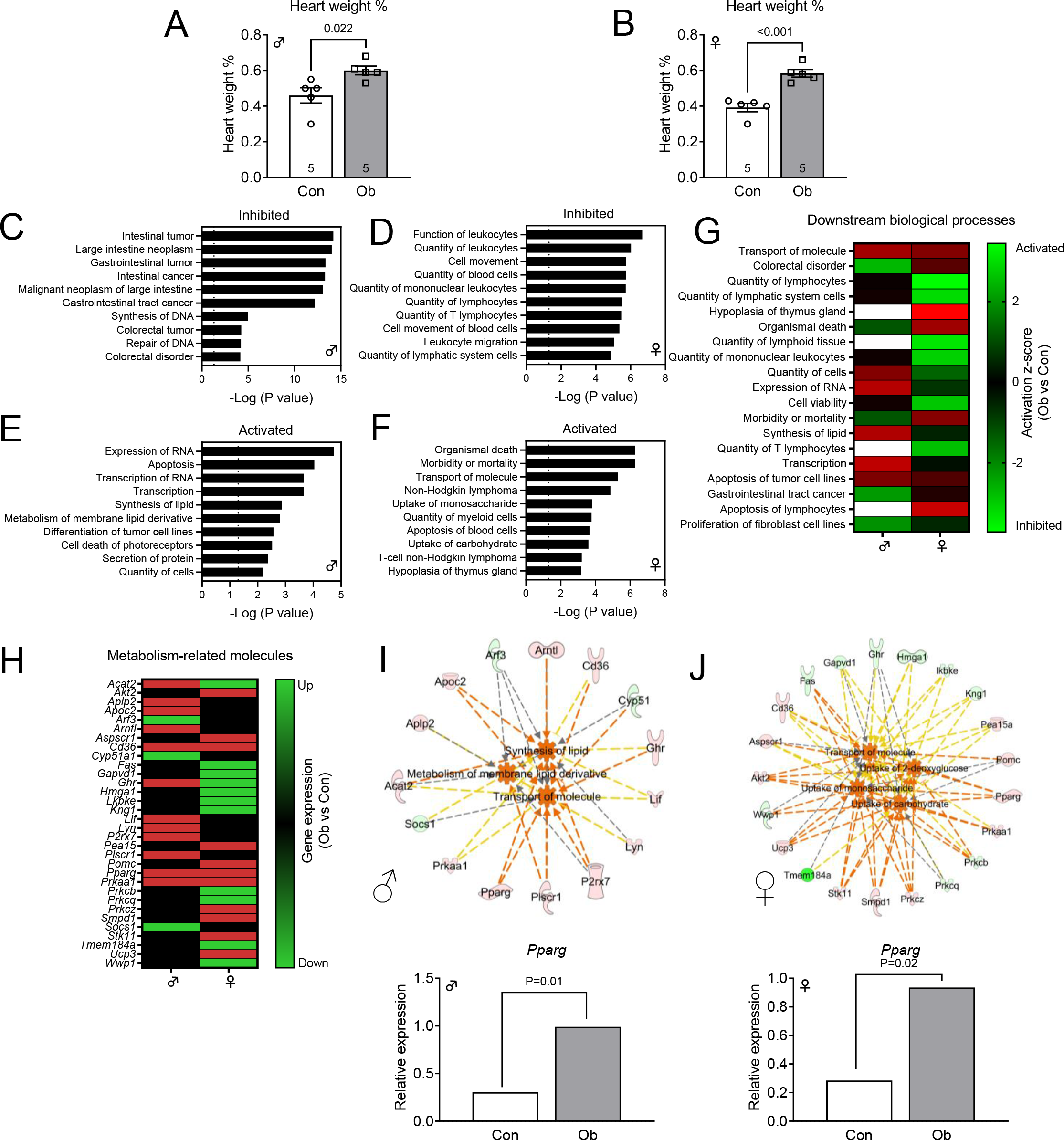
Maternal obesity induces fetal cardiac hypertrophy in association with altered transcription of metabolic genes at E18.5. (A, B) Heart weight, relative to body weight, in female and male fetuses of control (Con) and obese (Ob) dams, on E18.5 of gestation. Con and Ob groups compared by Student’s t-test. P and n values given in figure. Bars are mean ± SEM. Symbols represent mean values of all pups of each sex within one litter. (C – F) Top 10 biological processes, ranked by P value, predicted to be inhibited (C, D) or activated (E, F) in hearts of female and male fetuses of obese dams, based on Ingenuity Pathway Analysis of differential gene expression, compared to controls. Activation status determined by z-score >|1.7|. (G) Comparison of biological processes activated or inhibited in response to maternal obesity, in male and female fetuses. Color scale indicates activation z-score relative to control fetuses. (H) Comparison of metabolic genes differentially expressed in response to maternal obesity, in male and female fetuses. Color scale indicates differential expression relative to control fetuses. (I, J) Predicted effect of differentially expressed metabolic genes on downstream biological processes in hearts of female and male fetuses of obese dams. Color of molecules represents expression change Ob vs Con: red, upregulated; green, downregulated. Color of arrows represents direction of predicted effect on downstream processes: orange, activation; grey, neutral; yellow, expression inconsistent with predicted activation. (K, L) Least-squared mean relative PPARG expression in female and male fetal hearts from Con and Ob dams determined by RNASeq (n=5 of each sex per group). P values given in figure.

**Table 1.** Effect of maternal obesity on body and heart weights in E18.5 fetuses.

|  | Con | Ob | P value (t test) |
| --- | --- | --- | --- |
| ***Males*** | n=5 litters | n=5 litters |  |
| Body weight (g) | 1050 ± 50 | 1272 ± 26 | **0.004\*** |
| Heart weight (mg) | 4.84 ± 0.56 | 7.64 ± 0.28 | **0.002\*** |
| ***Females*** | n=5 litters | n=5 litters |  |
| Body weight (g) | 995 ± 43 | 1228 ± 58 | **0.012\*** |
| Heart weight (mg) | 3.88 ± 0.26 | 7.14 ± 0.32 | **<0.001\*** |

Body and heart weights, determined at necropsy, in male and female fetuses of control (Con) and obese (Ob) dams. Litter mean values for Con and Ob groups compared by Student's t-test. \*, P<0.05. Mean ± SEM.

Maternal obesity concomitantly altered the expression of 841 genes in the hearts of female fetuses and 764 genes in the hearts of male fetuses, with 66 genes commonly altered in both sexes (Suppl. Fig. 1A, B). Ingenuity analysis predicted inhibition of processes related to neoplasia and DNA repair/synthesis in male fetal hearts, and pathways related to function, quantity and movement of immune cells in female fetal hearts, in obese compared to control dams (z-score <-1.7, top ten functions by P-value, Fig. 1C, D). By contrast, maternal obesity activated synthesis of lipid and metabolism of membrane lipid derivatives in male fetuses, and uptake of monosaccharides and carbohydrates in female fetuses (activation z-score > 1.7, top-ten functions by P-value, Fig. 1 E, F). Transport of molecules was consistently activated by maternal obesity in both males and females (Fig. 1G).

The activated downstream functions related to molecular transport, carbohydrate and lipid metabolism encompassed an overlapping suite of differentially expressed genes, several of which were commonly regulated by maternal obesity in male and female fetuses (Fig. 1H). Specifically, maternal obesity upregulated *Pparg*, the nuclear peroxisome proliferator activated receptor implicated in lipogenesis, *Cd36,* a plasma membrane fatty acid translocase, and *Prkaa1*, the catalytic subunit of cytoplasmic AMP-activated protein kinase (AMPKa1), in fetuses of both sexes (Fig. 1H-J). Network analysis of direct molecular interactions between the differentially expressed, metabolism-related genes indicated that *Pparg* was a critical node in both the female and male fetal transcriptomic response (Suppl. Fig. 1C, D).

Furthermore, when unsupervised regulator effects analysis was used to link the annotated downstream functions of differentially expressed genes to upstream effectors, a network of genes activated by *Ppargc1a* and including *Cd36* and the triglyceride synthesis enzyme, *Lpin1*, was predicted to promote lipid synthesis in male fetal hearts in response to maternal obesity (Suppl. Fig 1E.). Taken together, the transcriptomic data were most consistent with maternal obesity promoting cardiac lipid metabolism in male and female fetuses, by increasing expression of genes related to fatty acid uptake, lipid synthesis and oxidation.

### Maternal obesity in pregnant mice causes age- and sex-dependent diastolic dysfunction and left ventricular dilation in adult offspring

In male offspring, E/A and E’/A’ ratios of left ventricular diastolic function declined overall with increasing postnatal age from 3 to 24 months and were further impaired by maternal obesity at all time points (Fig. 2A, C). Maternal obesity also reduced E/E’ ratio in males, with significant differences apparent between Con and Ob groups in both the youngest and oldest offspring studied (Fig. 2E). By contrast, the effect of maternal obesity on diastolic function in female offspring depended on the age at which they were studied. Three months after birth, neither E/A nor E’/A’ ratio differed between Con and Ob female offspring (Fig 2B, D). Six months after birth, E’/A’ ratio, but not E/A ratio, was reduced, and by 9 months both E’/A’ and E/A ratios were lower in Ob compared to Con females. Finally, at age 24 months, E/A ratio was higher in Ob than Con female offspring, whereas E’/A’ ratio was similar in the two groups, indicating a more severe state of diastolic dysfunction with pseudonormal filling pattern. E/E’ ratio was lower in Ob than Con female offspring, irrespective of age (Fig. 2F). Therefore, maternal obesity caused diastolic dysfunction in adult offspring of both sexes, albeit the effect was later in onset but faster in progression in females than males.

**Fig. 2.**
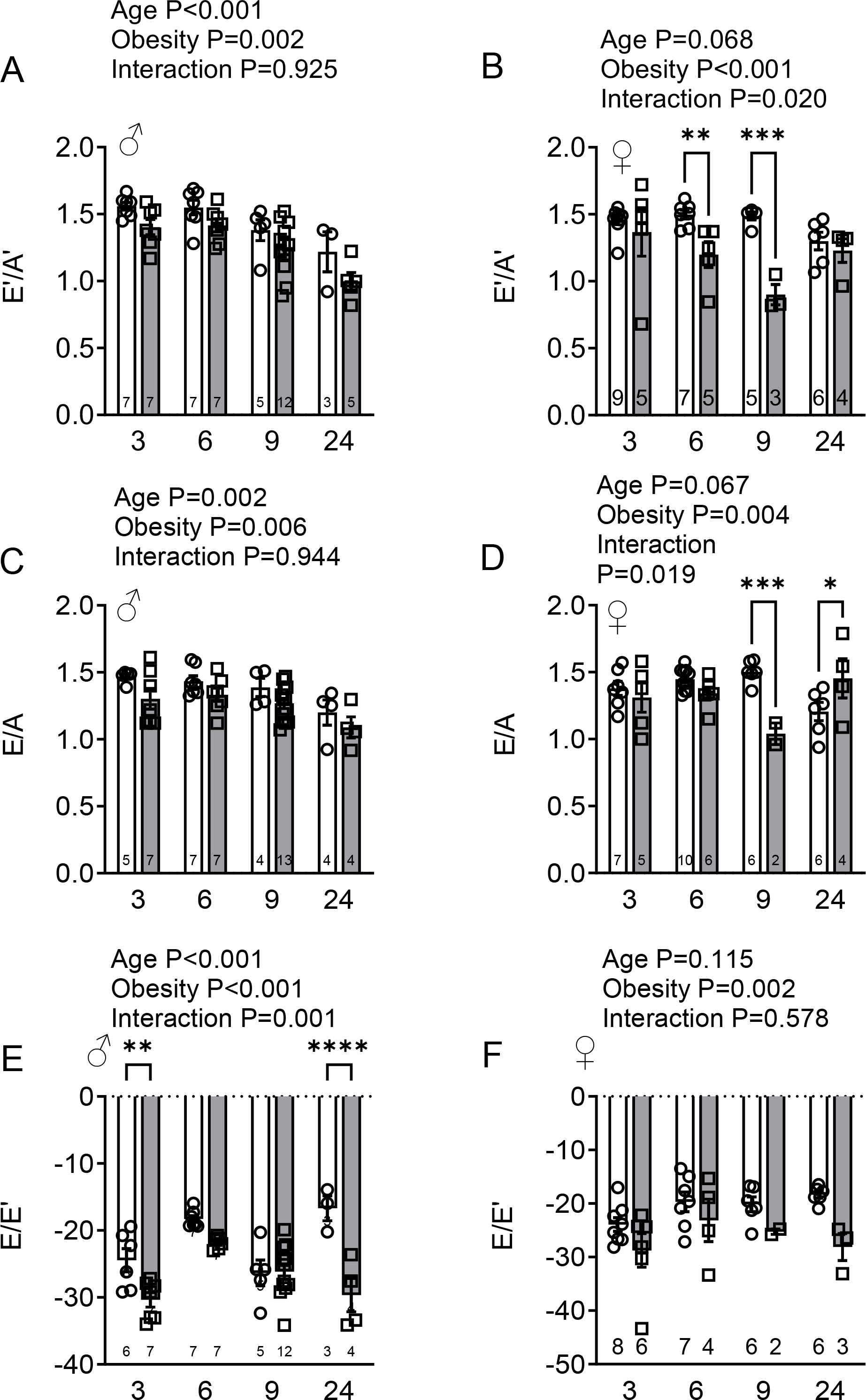
Maternal obesity causes age- and sex-dependent left ventricular diastolic dysfunction in adult offspring. Echocardiographic indices of diastolic function in 3- to 24-month old male (A, C, E) and female (B, D, F) offspring of control (white bars) and obese dams (grey bars). (A, B) Ratios of early- to late-diastolic left ventricular wall displacement determined by tissue Doppler (E’/A’). (C, D) Ratios of early- to late-diastolic mitral inflow determined by pulsed wave Doppler (E/A). (E, F) Ratio of early diastolic mitral inflow to wall displacement (E/E’). Main effects of maternal obesity and postnatal age, and their interaction, were determined by two-way ANOVA and P values given in figure. Post-hoc comparisons of Con and Ob groups at each age used the Sidak test; * P<0.05, ** P<0.01, ***P<0.001. Bars are mean ± SEM. Symbols represent individual animals. n values given in bars.

In males, maternal obesity increased end-diastolic volume in 2-year-old offspring and increased end-systolic volume irrespective of age (Fig. 3A, C). By contrast, in female offspring, maternal obesity increased left ventricular end-diastolic volume, but not end-systolic volume (Fig. 3B, D). Both left ventricular wall thickness and systolic function, measured by ejection fraction and fractional shortening, increased modestly with age in male and female mice but were not affected by maternal obesity (Suppl. Fig 2). When tissues were weighed at necropsy in 6-month old offspring, heart weight was significantly greater in Ob than Con females as an unadjusted value, but not expressed as a percentage of body weight (Table 2). However, none of the other measurements of body, heart or ventricle weight differed between Con and Ob offspring at this age.

**Fig. 3.**
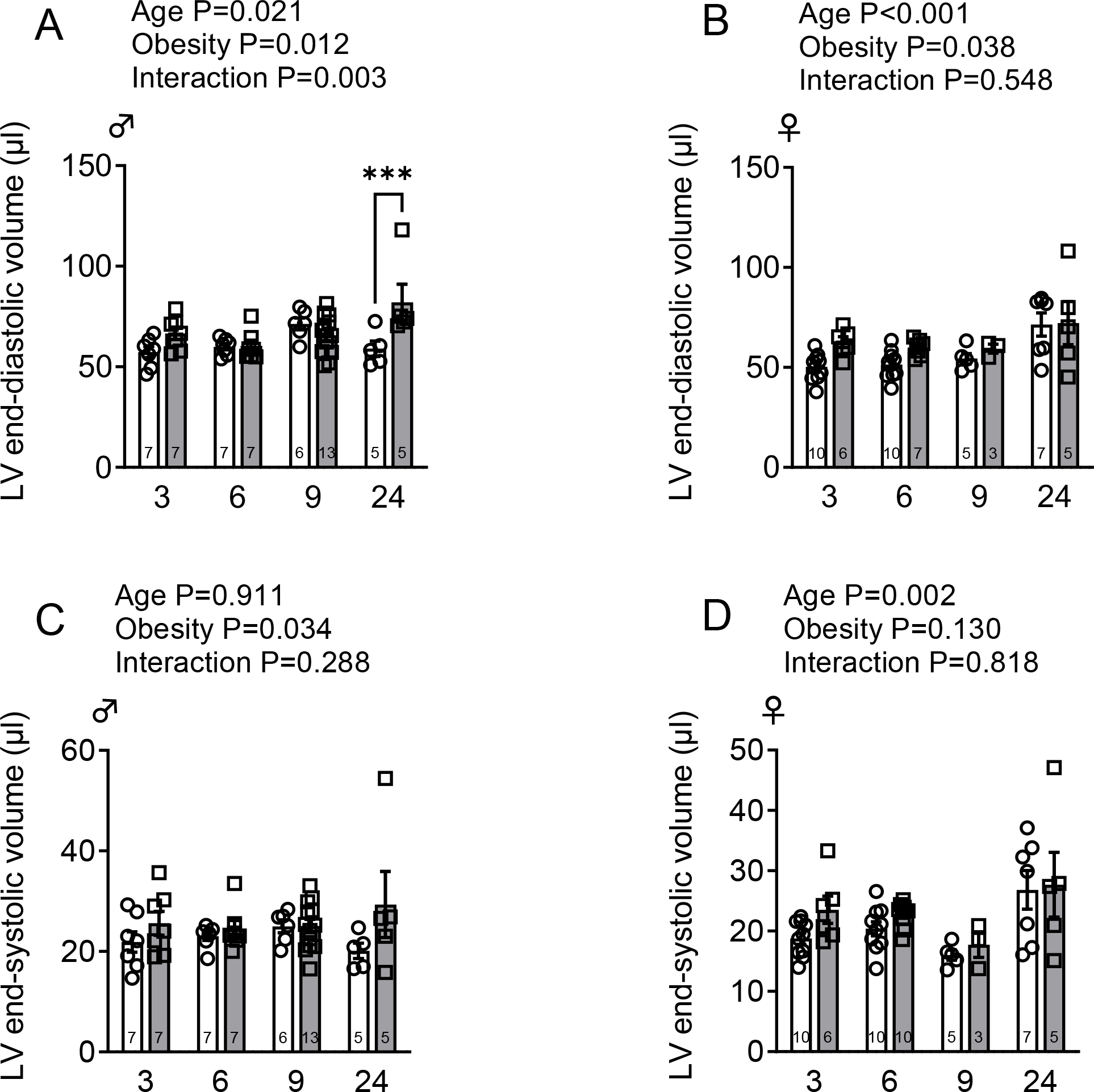
Maternal obesity causes age- and sex-dependent left ventricular dilatation in adult offspring. Left ventricular end-diastolic and end-systolic volumes, determined by echocardiography, in 3- to 24-month old male (A, C) and female (B, D) offspring of control (white bars) and obese dams (grey bars). Main effects of maternal obesity and postnatal age, and their interaction, were determined by two-way ANOVA and P values given in figure. Post-hoc comparisons of Con and Ob groups at each age used the Sidak test; * P<0.05, ** P<0.01, ***P<0.001 Bars are mean ± SEM. Points represent individual animals. n values given in figure.

**Table 2.** Effect of maternal obesity on body and heart weights in 6-month-old adult offspring.

|  | Con | Ob | P value (t test) |
| --- | --- | --- | --- |
| **Males** | n=47 | n=31 |  |
| Body weight (g) | 36.4 ± 0.9 | 36.1 ± 1.1 | 0.830 |
| Heart weight (mg) | 158 ± 3 | 162 ± 5 | 0.527 |
| (% body wt. x 1000) | 4.40 ± 0.10 | 4.55 ± 0.16 | 0.405 |
| LV weight (g) | 112 ± 2 | 114 ± 4 | 0.623 |
| (% body wt. x 1000) | 3.08 ± 0.07 | 3.24 ± 0.16 | 0.289 |
| **Females** | n=42 | n=34 |  |
| Body weight (g) | 26.8 ± 0.7 | 28.5 ± 0.9 | 0.149 |
| Heart weight (mg) | 127 ± 3 | 139 ± 4 | 0.012* |
| (% body wt. x 1000) | 4.79 ± 0.10 | 4.97 ± 0.16 | 0.305 |
| LV weight (g) | 89.7 ± 2.4 | 93.9 ± 3.54 | 0.325 |
| (% body wt. x 1000) | 3.35 ± 0.06 | 3.44 ± 0.13 | 0.561 |

Body, heart and LV weight, determined at necropsy, in 6-month old male and female offspring of control (Con) and obese (Ob) dams. Con and Ob groups compared by Student's t-test. \*, P<0.05. Mean ± SEM.

### Maternal obesity alters cardiac metabolism in adult offspring in a sex-specific manner

#### Gene expression

Maternal obesity upregulated *Pparg* expression in both male and female 6-month old offspring of obese dams, compared to controls (Fig. 4A, B). *Cd36* and *Prkaa1* expression were also higher in Ob than Con male offspring hearts, but not in females (Fig. 4). In male offspring, maternal obesity increased expression of downstream *Pparg* targets related to lipid synthesis and storage (*Fasn, Plin2, Srebp1)* and lipid oxidation (*Pgc1a, Cpt1b, Acox1*) but did not alter *Pgc1b, Cpt1a, Mcad, Hoad, Pdk4* or *Ucp3* expression (Fig. 4A). In female offspring, maternal obesity upregulated *Fasn* but did not affect expression of any of the other downstream mediators of lipid metabolism studied (Fig. 4B). When adult male offspring from control and obese dams were combined, there was a strong inverse correlation between cardiac *Pparg* gene expression and E/E’ ratio (Fig. 4C). In female offspring, there was a strong correlation between cardiac *Pparg* expression and left-ventricular end-diastolic volume (Fig. 4D). *Cd36* expression in female offspring also positively correlated with E/A ratio and inversely correlated with wall thickness at diastole, whilst both *Pdk4* and *Hoad* expression correlated with wall thickness at systole and end-diastolic volume (Suppl. Table 2). There were no other significant correlations between cardiac gene expression and the measured echocardiographic and morphometric outcomes in adult offspring (Suppl. Table 2). Therefore, transcriptional upregulation of *Pparg* expression and lipid metabolism persisted in the offspring of obese dams and was linked to cardiac functional and structural phenotype in male and female offspring.

**Fig. 4.**
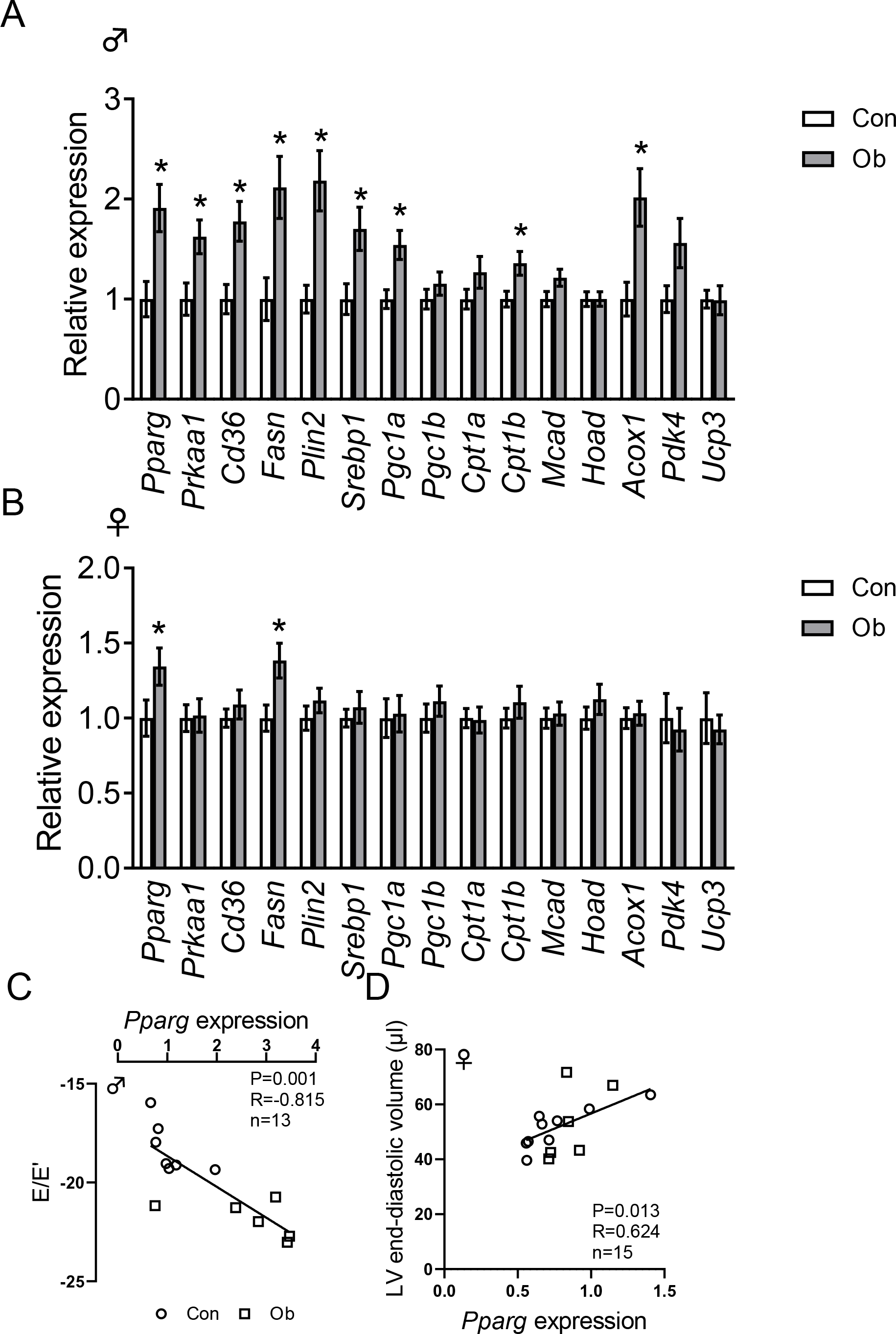
Maternal obesity increases cardiac expression of Pparg and its downstream targets in adult offspring. Relative expression of candidate genes in left ventricles of 6-month old old male (A, Con n=11, Ob n=10) and female (B, Con n=9, Ob n=10) offspring at age 6 months, determined by qPCR relative to *Rna18s* and *Rps29*. Bars are mean ± SEM. Con and Ob offspring compared by Student’s t-test.*, P<0.05. (C, D) Correlation of left ventricular *Pparg* expression with E/E’ ratio in male offspring and end-diastolic volume in female offspring of Con and Ob dams. Relationship between variables assessed by Pearson’s correlation, P, R and N values given in figure. Least-squares regression line shown.

#### Lipid metabolism

Left ventricular ^3^H-oleic acid clearance *in vivo* was greater in female than male offspring of control dams (Suppl. Fig. 3A). However, cardiac fatty acid clearance did not differ significantly between Ob offspring and their Con counterparts of the same sex (Fig. 5A, B). Maternal obesity reduced FATP6 transporter protein abundance in hearts of female, but not male offspring, and did not affect CD36 or FATP1 protein abundance (Table 3).

**Fig. 5.**
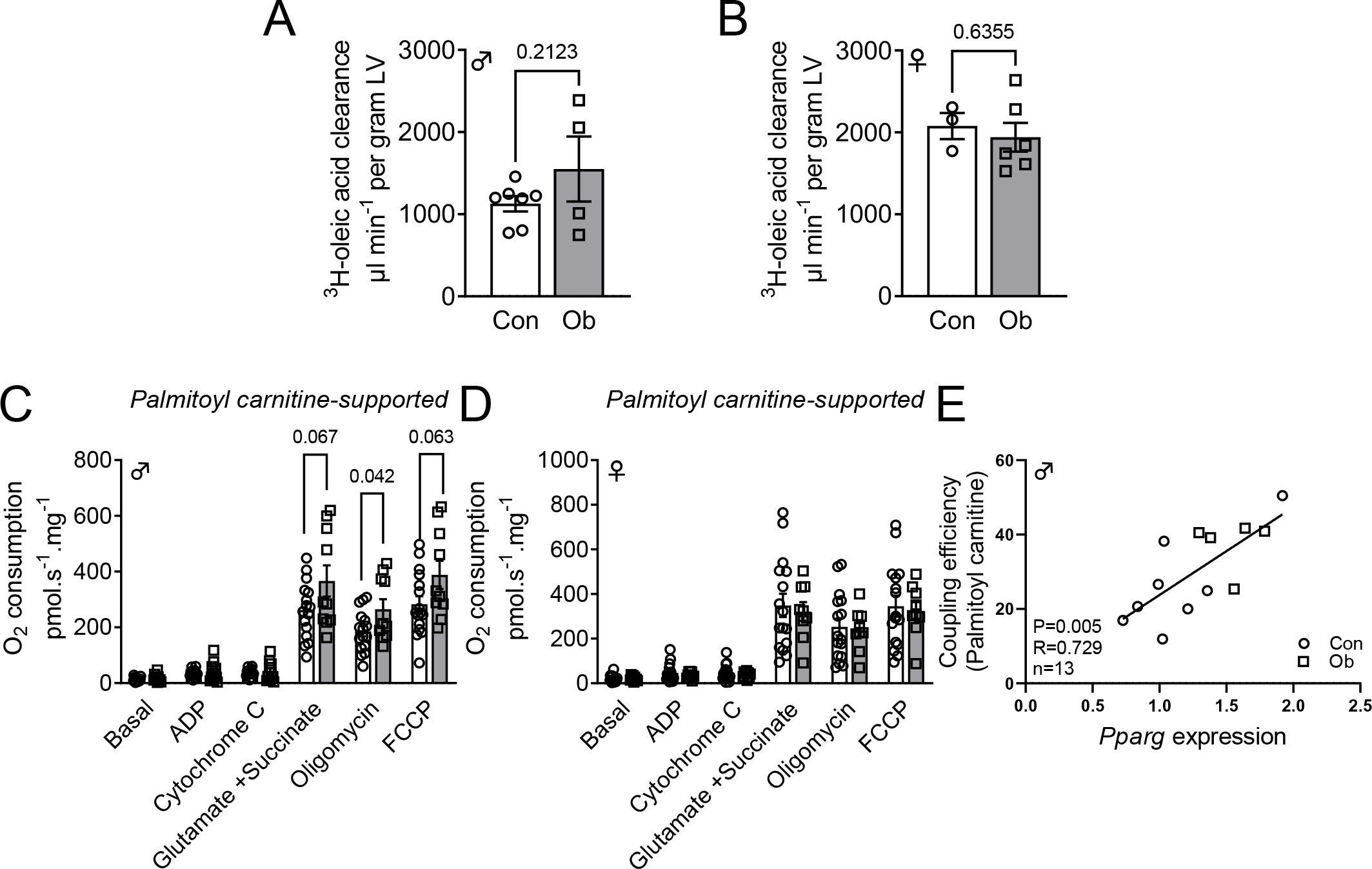
Maternal obesity increases myocardial fatty acid oxidation in adult male, but not female, offspring. (A, B) In vivo left ventricular 3H-oleic acid clearance in male (Con n=7, Ob n=4) and female (Con n=3, Ob n=6) offspring. (C, D) Palmitoyl carnitine-supported mitochondrial respiration rates in isolated, permeabilized cardiac myofibers from 6-month old male (Con n=16, Ob n=10) and female (Con n=16, Ob n=9) offspring of Con and Ob dams. Respiration rates and clearances compared between Con and Ob groups by Student’s t-test, P values given in figure. Mean ± SEM. Symbols represent individual offspring. (E) Correlation between palmitoyl carnitine supported coupling control ratio and Pparg expression in male offspring. Relationship between variables determined by Pearson’s correlation, P and R values in figure. Least-squares regression line shown.

**Table 3.** Effect of maternal obesity on cardiac fatty acid and glucose transporter expression in 6-month-old adult offspring.

|  | Con | Ob | P value (t test) |
| --- | --- | --- | --- |
| ***Males*** | n=10 | n=10 |  |
| CD36 | 1.00 ± 0.05 | 1.02 ± 0.06 | 0.748 |
| FATP1 | 1.00 ± 0.23 | 0.90 ± 0.32 | 0.331<sup>b</sup> |
| FATP6 | 1.00 ± 0.24 | 1.25 ± 0.28 | 0.501 |
| GLUT1 | 1.00 ± 0.10 | 1.18 ± 0.15 | 0.321 |
| GLUT4 | 1.00 ± 0.12 | 1.12 ± 0.11 | 0.448 |
| ***Females*** | n=10 | n=10 |  |
| CD36 | 1.00 ± 0.05 | 0.95 ± 0.06 | 0.462 |
| FATP1 | 1.00 ± 0.18 | 2.27 ± 0.67 | 0.143<sup>a</sup> |
| FATP6 | 1.00 ± 0.17 | 0.44 ± 0.06 | **0.006<sup>b\*</sup>** |
| GLUT1 | 1.00 ± 0.32 | 0.88 ± 0.34 | 0.310<sup>a</sup> |
| GLUT4 | 1.00 ± 0.03 | 1.02 ± 0.05 | 0.746 |

Relative transporter protein abundance in LV homogenates, determined by western blot, in 6-month old male and female offspring of control (Con) and obese (Ob) dams. Con and Ob groups compared by Student's t-test or Mann-Whitney test<sup>a</sup>, as appropriate to distribution. <sup>b</sup>Log transformed prior to statistical analysis. \*P<0.05. Mean ± SEM.

In male offspring, maternal obesity increased cardiac palmitoyl carnitine respiration in the leak state (+oligomycin), when palmitoyl carnitine was supplied in combination with malate, glutamate and succinate (Fig. 5C). Maximal oxidative phosphorylation-coupled palmitoyl carnitine respiration and electron transport chain capacity (+FCCP) also tended to be higher in Ob than Con male offspring hearts (Fig. 5C). However, maternal obesity had no effect on either coupled or leak respiration supported by palmitoyl carnitine alone in male offspring (Fig. 5C), or on any of the rates of fatty acid supported respiration in female offspring hearts (Fig. 5D). The coupling efficiency of fatty acid supported respiration correlated with cardiac *Pparg* expression in male offspring (Fig. 5E). However, there were no overall differences in coupling efficiency between Con and Ob offspring (Suppl. Table 3).

Maternal obesity did not alter adult offspring cardiac fatty acid content (Suppl. Table 4) or triglyceride content (females Con 7.7 ± 2.2 µg/mg, Ob 9.3 ± 2.2 µg/mg; males Con 22.9 ± 5.0 µg/mg, Ob 32.7 ± 6.2 µg/mg). Maternal obesity reduced offspring cardiac 1,2-18:2/18:1 DAG content but did not alter abundance of the other DAGs measured in adult male offspring and had no effect on any of the DAGs in female offspring hearts (Suppl. Table 5). There was no difference in ceramide content between Con and Ob offspring (Suppl. Table 6).

#### Carbohydrate metabolism

Myocardial glucose uptake, determined using PET as the rate of clearance of ^18^F-deoxyglucose from plasma into the left ventricle, was greater in male than female control offspring (Suppl. Fig. 3B, C). Maternal obesity reduced myocardial glucose uptake in female but not male offspring (Fig. 6A-D) but did not alter cardiac GLUT1 or GLUT4 abundance in offspring of either sex (Table 3). There were no differences between Con and Ob offspring in the rates of pyruvate-supported mitochondrial respiration in cardiac muscle *ex vivo* (Fig. 6E, F). Similarly, neither histone H3 protein abundance nor acetylation differed between Con and Ob offspring (Fig. 7).

**Fig. 6.**
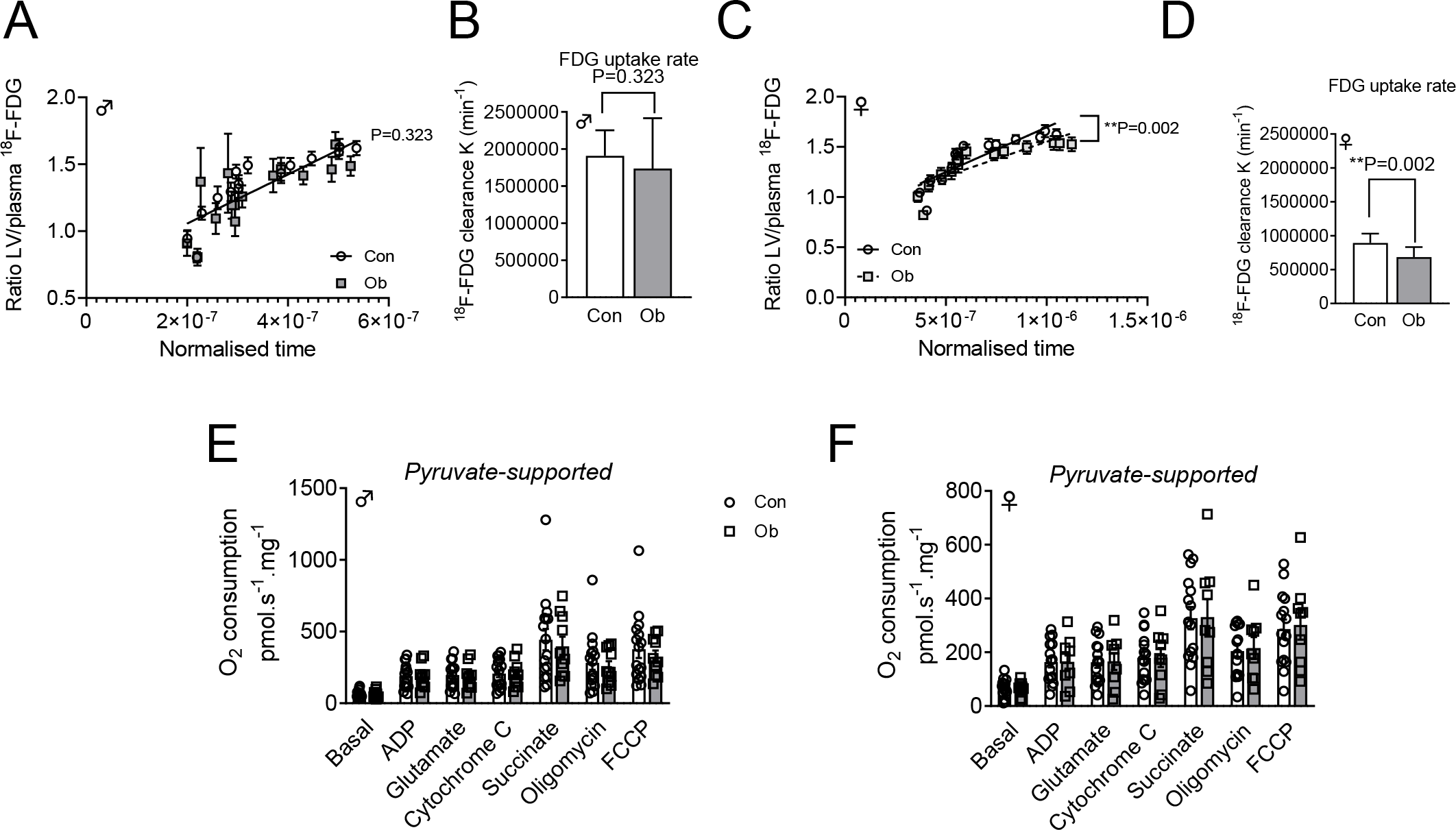
Maternal obesity impairs myocardial glucose uptake in adult female, but not male, offspring. (A-D) Left ventricular 8F-fluorodeoxyglucose clearance in male (Con n=8, Ob n=9) and female (Con n=10, Ob n=11) offspring of control and obese dams at age 6 months. Con and Ob groups compared by least-square linear regression of (A, C) Patlak plot, symbols represent mean ± SEM values for all individuals within each group, at each normalized time point. (B, D) Histogram of mean ± SEM clearance (gradient of Patlak plot) for each experimental group. (E, F) Pyruvate-supported mitochondrial respiration rates in isolated, permeabilized cardiac myofibers from 6-month old male (Con n=16, Ob n=10) and female (Con n=15, Ob n=9) offspring of control and obese dams. Con and Ob groups compared by Student’s t test. P values for intergroup comparisons given in figure. Bars are mean + SEM. Points represent individual animals.

**Fig. 7.**
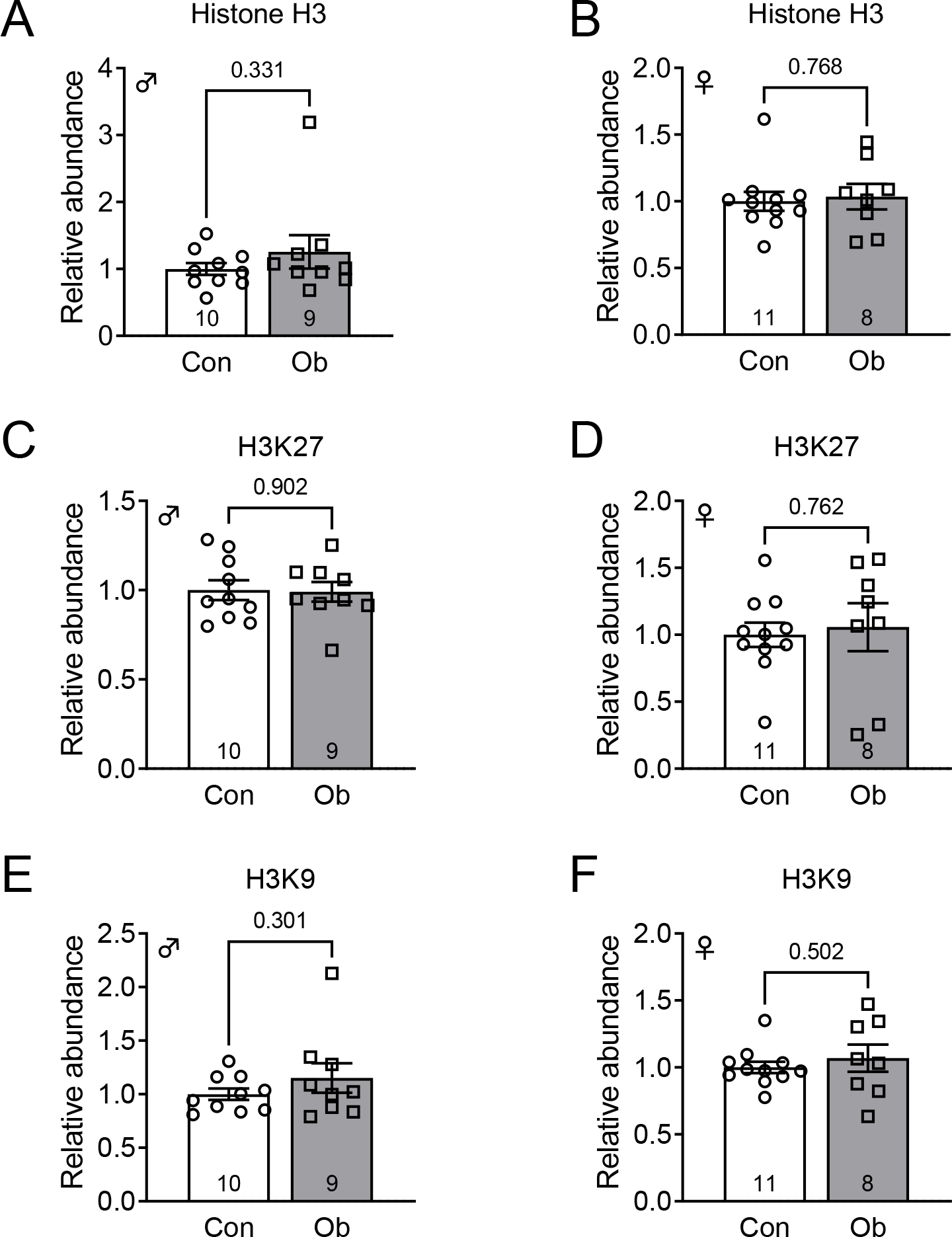
Maternal obesity does not affect myocardial histone H3 acetylation in adult offspring. Relative protein abundance of total (A, B), acetyl-lysine-27 (C, D) and acetyl-lysine-9 (E, F) histone H3 in 6-month old male and female offspring of Con and Ob dams. Con and Ob groups compared by Student’s t-test, P values given in figure. Bars are mean ± SEM. Points represent individual animals. n values given in figure.

## DISCUSSION

This is the first study in mice to link the effects of maternal obesity on fetal cardiac morphology and gene expression with alterations in cardiac contractile and metabolic function in the adult offspring. The results show that maternal obesity causes fetal cardiac hypertrophy *in utero*, in association with transcriptomic changes consistent with altered cardiac carbohydrate and lipid metabolism. We also report that maternal obesity impairs cardiac diastolic function in both male and female adult offspring, up to 2 years after birth, but the progression of cardiac dysfunction with age is sex dependent. The impairments in contractile function were accompanied by persistent *Pparg* upregulation and altered carbohydrate and lipid metabolism in female and male offspring, respectively. The study therefore indicates that cardiac metabolism is programmed by *in utero* exposure to maternal obesity and contributes to cardiac dysfunction in later life.

The hypertrophic effects of maternal obesity on fetal heart and left ventricle weight were similar in female and male fetuses and consistent with that reported previously in other experimental animals ^9–12^ and pregnant women ^6, 7^. In contrast, the effect on the fetal cardiac transcriptome strongly depended on fetal sex, with less than 10% of the differentially expressed genes shared between females and males. Based on bioinformatic analysis of the differentially expressed genes, the gross hypertrophic effect did not appear to be explained by cellular processes that were transcriptionally inhibited in response to maternal obesity, which were broadly related to inflammation in female fetuses and cell proliferation in male fetuses. These changes could be linked to myocardial fibrotic remodelling ^10, 11^ or impaired cardiomyocyte endowment, depending on fetal sex. However, the processes that were transcriptionally activated seemed more likely to contribute to cardiac hypertrophy, because they were associated with transport, uptake, synthesis and metabolism of macronutrient molecules required for cell growth and contractile function. Certainly, cardiac lipid and glucose metabolism are strongly linked to cardiac dysfunction in obese and diabetic adults ^19, 20^.

Our bioinformatic analysis pointed to *Pparg* as a central node in the network of differentially expressed genes related to metabolism. Cardiac *Pparg* was upregulated in both female and male fetuses of obese dams, consistent with previous reports in liver and skeletal muscle from fetuses of pregnant macaques fed a high fat diet ^30, 31^. The PPARγ protein is a transcription factor that is activated by binding with fatty acid ligands and drives the expression of genes mediating fatty acid uptake and oxidation, including fatty acid translocase *Cd36*, which was also upregulated in fetuses of obese dams. *Pparg* mRNA expression is upregulated in adult mice fed a high fat diet ^32^ and cardiomyocyte-specific overexpression of *Pparg* causes cardiac hypertrophy in transgenic adult mice, in association with systolic dysfunction, abnormal mitochondrial architecture, increased triglyceride uptake and lipid storage and increased expression of genes encoding for proteins involved in β-oxidation ^21^. Our previous study showed that maternal obesity increases placental lipid transport and fetal lipid load in mice ^33^. Therefore, fetal cardiac hypertrophy in pregnancies complicated by maternal obesity may be promoted by increased circulating fatty acid availability activating PPARγ signalling in the heart.

The finding that maternal obesity impairs diastolic function in adult offspring up to 2 years old extends our previous observations in this model ^23^. These data confirm that transient cardiac hypertrophy due to excess nutrition in early life leads to lasting impairments in contractile function of the heart, in common with other studies in mice and their isolated cardiomyocytes ^5, 18^. Since systolic function was not affected by maternal obesity, the echocardiographic observations in offspring of obese dams were most consistent with the phenotype of heart failure with preserved ejection fraction, which is a significant cause of cardiovascular mortality in people ^34^. Mild diastolic dysfunction is often subclinical and not diagnosed as heart failure, but still associated with increased mortality. Our data therefore suggest diastolic dysfunction or heart failure could be a contributing factor to increased later life cardiovascular morbidity in people whose mothers had obesity during pregnancy ^3^. To our knowledge, there have been no studies of adult cardiac function in this population.

In contrast with the changes in fetal heart weight, the effect of maternal obesity on offspring cardiac function depended on postnatal age and sex, with diastolic function consistently impaired in males but progressively worsening with age in females. This may be partly explained by circulating oestrogens having a cardioprotective effect in young female offspring ^35^. The appearance of overt diastolic dysfunction in 9-month old offspring of obese dams, with both E’/A’ and E/A ratios reduced, approximately coincided with declining oestradiol levels in this strain and may therefore reflect diminishment of the protective oestrogen effect ^36^. Unfortunately, we did not consider the influence of reproductive cycles in the timing of our analyses and blood collections, preventing us from further investigating the contribution of oestradiol levels. Female offspring of obese dams had most severe diastolic dysfunction at 2 years of age, which is certainly after full reproductive senescence occurs between 13 and 16 months ^37^. Sex differences in the long-term effect of maternal obesity on the offspring heart could therefore be due to sexually dimorphic changes in normal physiology and endocrinology with age. This is consistent with clinical observations showing that elderly women have a higher risk of cardiac dysfunction than men and are more likely to develop dysfunction in association with type 2 diabetes and ventricular hypertrophy ^38^.

Our study suggests that the sex-specific effects of maternal obesity may also be linked to differences in the metabolism of the heart. The results provide the first demonstration that *in vivo* cardiac metabolism depends on sex in adult mice, with greater cardiac fatty acid uptake in females but greater glucose uptake in males, in line with the known differences between women and men ^39, 40^. Reduced *in vivo* cardiac glucose uptake in adult female offspring of obese dams is consistent with the reported reduction in *in vitro* cardiomyocyte glucose uptake when the offspring of obese mice are fed a high fat diet ^18^. It also reflects the pathophysiological changes in the hearts of people with type 2 diabetes ^19^. Since there was no accompanying change in cardiac mitochondrial capacity for carbohydrate respiration or myocardial glucose transporter abundance, reduced cardiac glucose uptake may be explained by myocardial insulin resistance in female offspring of obese dams. Maternal obesity has been shown to cause peripheral insulin resistance, reduced cardiac insulin receptor abundance and sex-specific alterations in signalling pathways downstream of the insulin receptor in the offspring heart ^4, 23^. In turn, impaired glucose uptake may have limited cardiac flexibility to generate ATP under conditions of high workload or low oxygen, causing contractile dysfunction. Certainly, glucose uptake is positively correlated with systolic function in humans^41^.

Alterations in net cardiac lipid uptake did not explain diastolic dysfunction in the offspring of obese dams, even though myocardial fatty acid uptake has been reported to increase in association with diastolic dysfunction in people with diabetes ^19^. Cardiac dysfunction in adult male offspring in our study may instead have been related to the increase in mitochondrial fatty acid oxidation, which similarly occurs in association with reduced energetic efficiency in the hearts of adult rats fed a high fat diet ^42^. This effect may partly be mediated by increased expression of PPARγ co-activator 1α (*Pgc1a*), which stimulates mitochondrial biogenesis, and the carnitine acetyltransferase *Cpt1b*, responsible for trafficking long chain fatty acids into the mitochondria. Although our observations indicated that maternal obesity most robustly increased oligomycin-uncoupled β-oxidation, there was no accompanying increase in *Ucp3* expression or coupling control ratio, suggesting that the effect of maternal obesity was primarily due to increased activity of the β-oxidation pathway itself. Increased lipid oxidation in offspring of obese dams may have caused cardiomyocyte damage by increasing production of reactive oxygen species ^17^. Despite increased *Plin2* and *Srebp1* expression in males, we did not find evidence of increased cardiac lipid storage or altered abundance of DAG and ceramide derivatives, arguing against a role for cardiac lipotoxicity *per se* in the offspring of obese dams. This finding contrasted with previous studies in sheep fetuses and neonatal rats showing that maternal obesity increases cardiac lipid accumulation in the perinatal period ^9, 17^. Increased lipid oxidation may therefore outweigh increased storage in the long term. Indeed, cardiac triglyceride content is reduced in adult offspring of pigs fed a high fat diet ^43^. The results are therefore most consistent with increased lipid oxidation underpinning cardiac dysfunction in male offspring of obese dams.

At a molecular level, increased cardiac lipid oxidation was consistent with the persistant upregulation of *Pparg* expression, compared to controls. We therefore speculate that *Pparg* upregulation causes the myocardial metabolic alterations and, in turn, diastolic dysfunction in the adult male offspring. This proposal is supported by the strong correlation of *Pparg* expression with E/E’ ratio and respiratory coupling control ratio in adult male offspring. *Pparg* expression is strongly linked to increased locus specific H3K9 and H3K27 acetylation during adipogenesis ^44^ and maternal high fat feeding increases fetal hepatic *Pparg* expression in association with increased histone acetylation ^45^ but we did not find any changes in total cardiac H3K9 and H3K27 acetylation in this study. Therefore, the epigenetic mechanism underpinning persisting upregulation of *Pparg* in the offspring of obese dams remains unclear.

Taken together, the results support our hypothesis that maternal obesity alters cardiac metabolism in fetuses and adult offspring of obese pregnant mice. They show that the long-term effects are sex-specific but associated with cardiac *Pparg* upregulation and a shift from glucose to lipid metabolism in both male and female offspring of obese dams. Reduced cardiac metabolic flexibility, established *in utero*, may therefore contribute to later-life cardiometabolic disease risk in children of women with obesity. The findings imply that therapeutic strategies that modify the supply of nutrients to fetuses of obese women may improve their later health.

## SUPPLEMENTARY MATERIAL

### SUPPLEMENTARY FIGURE LEGENDS

**Suppl. Fig. 1.**
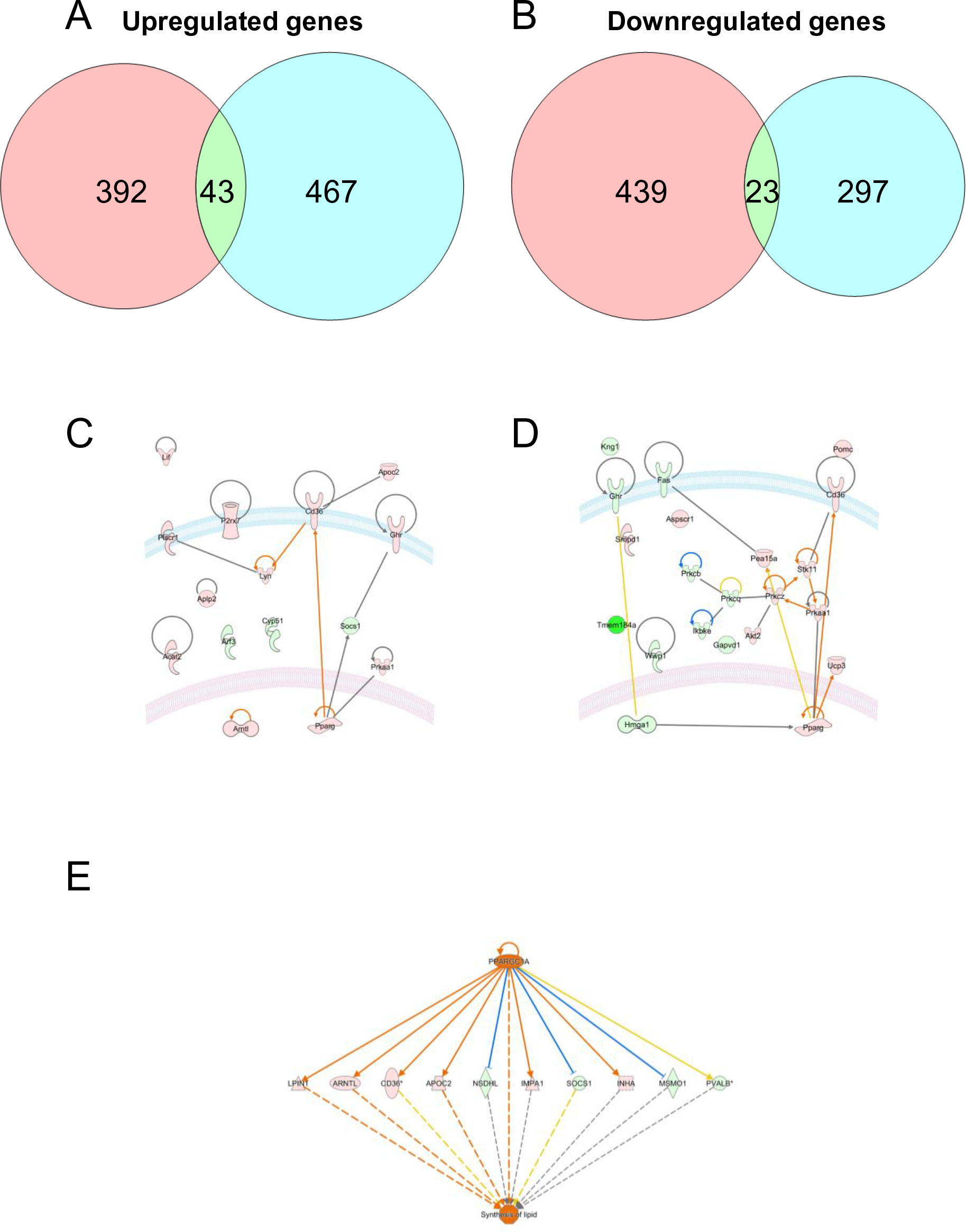
*Pparg* is a critical node in the transcriptional effects of maternal obesity on the fetal heart. (A, B) Venn diagrams to show numbers and intersections of up- and down-regulated genes in male and female fetuses. (C, D) Cellular location and direct molecular interactions of differentially expressed, metabolism-related genes in E18.5 male and female fetuses of obese dams. (E) Regulator effects network identified from differentially expressed genes in male fetuses of obese dams. RNASeq data annotated and figures generated by Ingenuity Pathway Analysis. Red symbols indicate upregulated genes, green symbols indicate downregulated genes. Orange arrows indicate predicted activation, blue arrows indicate predicted inhibition.

**Suppl. Fig. 2.**
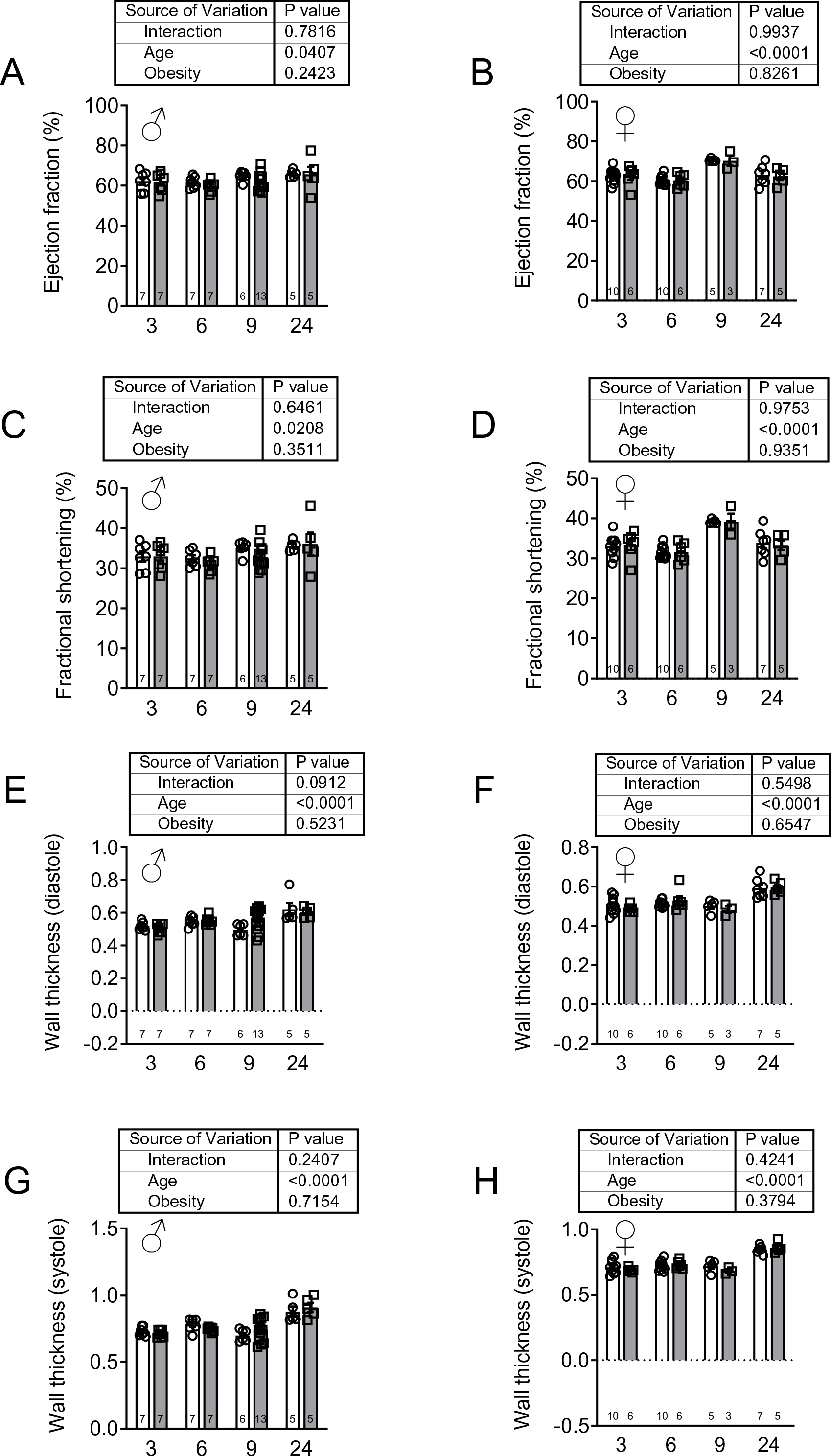
Maternal obesity does not alter left ventricular systolic function or wall thickness in adult offspring. (A, B) Ejection fraction, (C,D) fractional shortening, (E, F) wall thickness in diastole and (G, H) wall thickness at systole in male (A, C, E, G) and female (B, D, F, H) offspring of control and obese dams at 3, 6, 9 and 24 months after birth. Main effects of maternal obesity and postnatal age, and their interaction, were determined by two-way ANOVA and P values given in figure. Bars are mean ± SEM. Symbols represent individual animals. n values given in bars.

**Suppl. Fig. 3.**
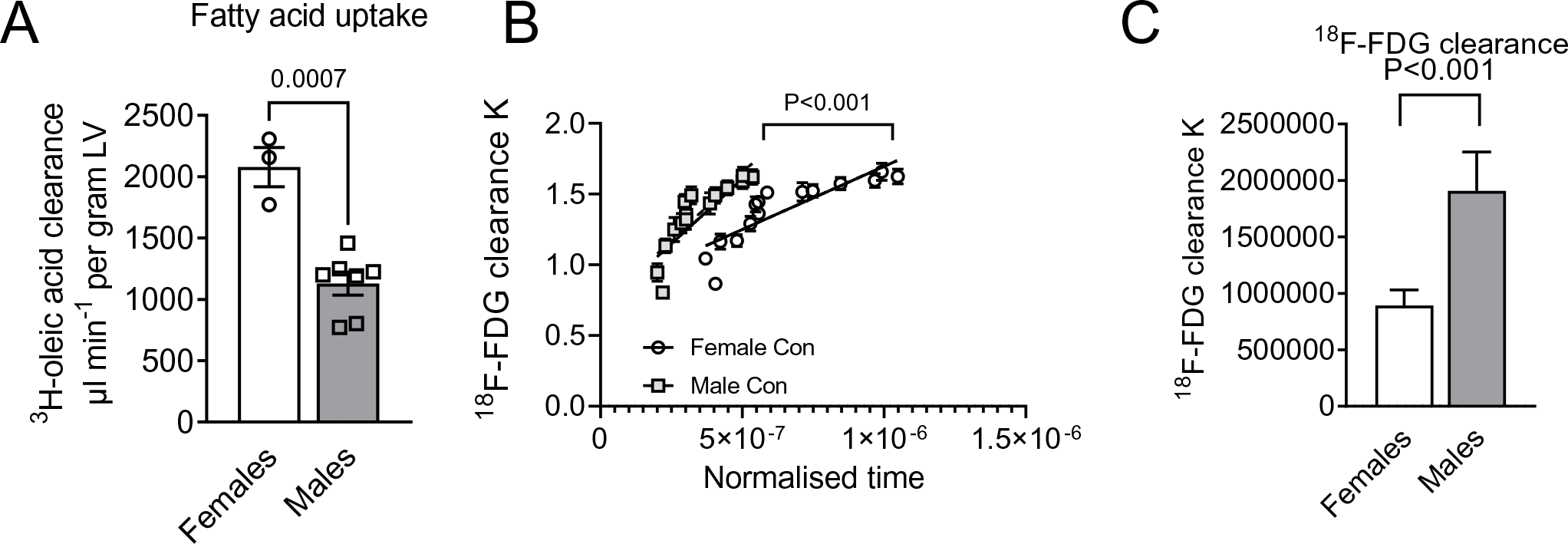
Cardiac *in vivo* fatty acid and glucose uptake differ between female and male offspring of control dams. (A) Left ventricular 3H-oleic acid clearance in female (Con n=3) and male (Con n=7) offspring. Con and Ob groups by Student’s t-test, Symbols represent individual offspring. (B,C) Left ventricular 8F-fluorodeoxyglucose clearance in female (Con n=10) and male (Con n=8) offspring of control and obese dams (B) Patlak plot, symbols represent mean ± SEM values for all individuals within each group, at each normalized time point. (C) Histogram of mean ± SEM clearance (gradient of Patlak plot) for each experimental group. P values given in figure. Mean ± SEM.

### SUPPLEMENTARY TABLES

**Suppl. Table 1.**
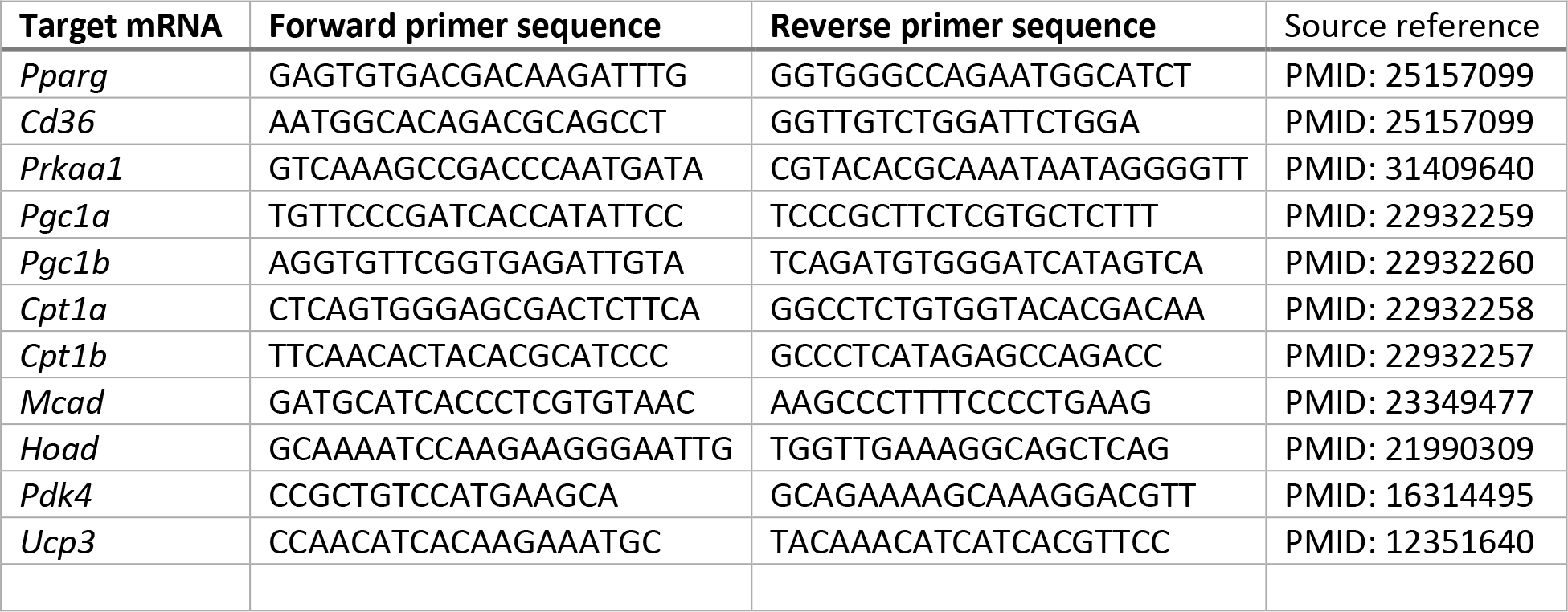
Primer sequences used for qRT-PCR analysis of adult offspring hearts.

**Suppl. Table 3.** Effect of maternal obesity on mitochondrial coupling efficiency in adult offspring.

|  | Con | Ob | P value (t test) |
| --- | --- | --- | --- |
| ***Males*** | n=16 | n=10 |  |
| Palmitoyl-carnitine supported respiration | 33.4 ± 2.0 | 31.5 ± 3.6 | 0.625 |
| Pyruvate-supported respiration | 26.8 ± 7.5 | 23.2 ± 6.1 | 0.218 |
| ***Females*** | n=15 | n=9 |  |
| Palmitoyl-carnitine supported respiration | 29.0 ± 1.6 | 27.0 ± 2.3 | 0.475 |
| Pyruvate-supported respiration | 28.4 ± 8.0 | 28.9 ± 8.1 | 0.871 |

Coupling efficiencies for each of the two combinations of substrates was calculated as 1 minus the ratio (leak state respiration/maximal electron transport chain capacity).

**Suppl. Table 4.** Effect of maternal obesity on adult offspring myocardial fatty acid content.

| nmol/mg protein | Con | Ob | P value (t-test) |
| --- | --- | --- | --- |
| **Males** | n=10 | n=10 |  |
| 12:0 | 17.8 ± 8.7 | 6.9 ± 4.7 | 0.991 |
| 14:0 | 50.1 ± 28.3 | 55.0 ± 31.0 | 1.000 |
| 16:0 | 1983.0 ± 424.3 | 1984.0 ± 432.4 | 1.000 |
| 16:1 n-7 | 16.6 ± 1.5 | 17.3 ± 6.0 | 1.000 |
| 18:0 | 1970.0 ± 264.7 | 2025.0 ± 227.0 | 1.000 |
| 18:1 n-9 | 1095.0 ± 121.6 | 1037.0 ± 189.8 | 1.000 |
| 18:2 n-6 | 4971.0 ± 615.8 | 4790.0 ± 405.0 | 1.000 |
| 18:3 n-3 | 7.3 ± 0.7 | 6.6 ± 0.5 | 0.999 |
| 18:3 n-6 | 0.01 ± 0.01 | 0.01 ± 0.004 | 1.000 |
| 20:1 n-9 | 62.7 ± 6.3 | 63.3 ± 3.4 | 1.000 |
| 20:3 n-6 | 73.2 ± 6.6 | 66.9 ± 3.6 | 0.999 |
| 20:4 n-6 | 1666.0 ± 148.6 | 1745.0 ± 79.8 | 1.000 |
| 20:5 n-3 | 8.9 ± 10.1 | 15.5 ± 12.3 | 1.000 |
| 22:4 n-6 | 81.9 ± 8.0 | 59.9 ± 6.1 | 0.496 |
| 22:5 n-3 | 255.8 ± 28.5 | 185.3 ± 12.4 | 0.460 |
| 22:6 n-3 | 2927.0 ± 224.4 | 2547.0 ± 192.4 | 0.973 |
| 22:5 n-6 | 270.7 ± 18.8 | 255.3 ± 10.2 | 0.999 |
| Total fatty acids | 15463 ± 1554 | 14859 ± 1288 | 0.768 |
| **Females** | n=10 | n=10 |  |
| 12:0 | 14.2 ± 9.1 | 0.9 ± 4.1 | 0.981 |
| 14:0 | 91.4 ± 57.3 | 3.8 ± 14.4 | 0.875 |
| 16:0 | 2183.0 ± 709.7 | 1064.0 ± 244.6 | 0.113 |
| 16:1 n-7 | 15.8 ± 5.1 | 14.1 ± 2.6 | 0.996 |
| 18:0 | 2394 ± 350.3 | 1817.0 ± 148.6 | 0.644 |
| 18:1 n-9 | 1181.0 ± 358.8 | 746.6 ± 155.3 | 0.595 |
| 18:2 n-6 | 5859.0 ± 391.5 | 5489.0 ± 423.3 | 0.195 |
| 18:3 n-3 | 9.9 ± 1.4 | 8.9 ± 0.8 | 0.999 |
| 18:3 n-6 | 27.4 ± 18.1 | 17.4 ± 22.7 | 0.985 |
| 20:1 n-9 | 55.7 ± 6.0 | 38.8 ± 3.9 | 0.982 |
| 20:3 n-6 | 54.0 ± 2.2 | 48.1 ± 3.6 | 1.000 |
| 20:4 n-6 | 1936.0 ± 99.2 | 1679.0 ± 87.3 | 0.998 |
| 20:5 n-3 | 20.5 ± 19.6 | -14.7 ± 9.4 | 0.929 |
| 22:4 n-6 | 68.2 ± 3.3 | 65.9 ± 5.3 | 0.991 |
| 22:5 n-3 | 138.6 ± 9.9 | 131.8 ± 8.5 | 0.986 |
| 22:6 n-3 | 3008.0 ± 226.9 | 2732.0 ± 204.2 | 0.850 |
| 22:5 n-6 | 213.2 ± 11.2 | 212.9 ± 17.5 | 0.947 |
| Total fatty acids | 17269 ± 1641 | 14055 ± 1083 | 0.120 |

Fatty acid content, determined by GC/MS, in LV homogenates from 6-month old male and female offspring of control (Con) and obese (Ob) dams. Con and Ob groups compared by Student's t-test with Holm-Sidak multiplicity correction. \*P<0.05. Mean ± SEM.

**Suppl. Table 5.** Effect of maternal obesity on adult offspring myocardial DAG content.

| pmol/mg protein | Con | Ob | P value (t-test) |
| --- | --- | --- | --- |
| **Males** | n=6 | n=5 |  |
| 1,2-16:0/18:0DG | 7.1 ± 2.1 | 5.1 ± 1.2 | 0.997 |
| 1,2-16:0/18:1DG | 34.3 ± 13.8 | 12.2 ± 3.3 | 0.434 |
| 1,2-16:0/18:2DG | 22.4 ± 9.3 | 8.3 ± 3.4 | 0.881 |
| 1,2-16:0/20:4DG | 4.4 ± 2.1 | 1.8 ± 1.5 | 0.997 |
| 1,2-1,2-16:0/22:6DG | 11.7 ± 3.9 | 5.3 ± 2.9 | 0.997 |
| 1,2-18:0/18:1DG | 9.4 ± 4.0 | 2.2 ± 0.8 | 0.997 |
| 1,2-18:0/18:2DG | 12.7 ± 4.0 | 6.8 ± 1.8 | 0.997 |
| 1,2-18:0/20:4DG | 55.1 ± 12.8 | 25.9 ± 6.3 | 0.128 |
| 1,2-18:0/22:6DG | 9.7 ± 3.3 | 3.9 ± 1.4 | 0.997 |
| 1,2-18:2/18:1DG | **52.3 ± 21.9** | **17.5 ± 4.7** | 0.033 |
| 1,2-di16:0DGa | 12.3 ± 3.5 | 5.4 ± 0.9 | 0.997 |
| 1,2-di18:0DGa | 5.2 ± 2.4 | 0.8 ± 0.5 | 0.997 |
| 1,2-di18:1DGa | 34.4 ± 16.2 | 7.0 ± 2.3 | 0.178 |
| 1,2-di18:2DGa | 28.0 ± 9.7 | 12.0 ± 4.1 | 0.816 |
| Total DAGs | 305 ± 105 | 115 ± 32 | 0.084<sup>b</sup> |
| **Females** | n=8 | n=6 |  |
| 1,2-16:0/18:0DG | 9.7 ± 1.6 | 9.8 ± 1.8 | 1.000 |
| 1,2-16:0/18:1DG | 35.4 ± 7.1 | 29.6 ± 4.7 | 0.997 |
| 1,2-16:0/18:2DG | 23.7 ± 5.4 | 19.0 ± 3.1 | 0.997 |
| 1,2-16:0/20:4DG | 5.5 ± 1.7 | 4.9 ± 1.4 | 1.000 |
| 1,2-1,2-16:0/22:6DG | 15.2 ± 3.5 | 10.5 ± 3.1 | 0.997 |
| 1,2-18:0/18:1DG | 11.5 ± 2.6 | 8.9 ± 1.7 | 0.999 |
| 1,2-18:0/18:2DG | 21.9 ± 3.2 | 16.8 ± 3.2 | 0.997 |
| 1,2-18:0/20:4DG | 56.5 ± 7.0 | 41.1 ± 9.3 | 0.421 |
| 1,2-18:0/22:6DG | 18.8 ± 4.8 | 13.1 ± 4.4 | 0.997 |
| 1,2-18:2/18:1DG | 58.0 ± 10.5 | 38.9 ± 5.5 | 0.147 |
| 1,2-di16:0DGa | 13.5 ± 2.1 | 12.0 ± 1.4 | 0.999 |
| 1,2-di18:0DGa | 5.2 ± 0.6 | 5.0 ± 1.3 | 1.000 |
| 1,2-di18:1DGa | 31.4 ± 8.1 | 22.8 ± 3.9 | 0.958 |
| 1,2-di18:2DGa | 42.5 ± 8.5 | 27.7 ± 6.0 | 0.456 |
| Total DAGs | 353 ± 67 | 261 ± 47 | 0.313 |

DAG content, determined by LC/MS, in LV homogenates from 6-month old male and female offspring of control (Con) and obese (Ob) dams. Con and Ob groups compared by Student's t-test with Holm-Sidak multiplicity correction. <sup>a</sup>Log transformed prior to statistical analysis. \*, P<0.05. Mean ± SEM.

**Suppl. Table 6.** Effect of maternal obesity on adult offspring myocardial ceramide content.

| pmol/mg protein | Con | Ob | P value (t-test) |
| --- | --- | --- | --- |
| **Males** | n=6 | n=5 |  |
| 16:0Cer | 7.8 ± 2.2 | 3.5 ± 1.0 | 0.852 |
| 18:0Cer | 5.1 ± 1.4 | 2.3 ± 0.6 | 0.852 |
| 20:0Cer | 13.5 ± 5.3 | 7.0 ± 1.9 | 0.390 |
| 22:0Cer | 16.6 ± 4.6 | 8.2 ± 1.8 | 0.181 |
| 24:0Cer | 5.5 ± 1.3 | 3.8 ± 0.7 | 0.852 |
| 24:1Cer | 8.1 ± 1.9 | 4.2 ± 1.1 | 0.777 |
| Total ceramides | 56.6 ± 16.4 | 29.0 ± 6.8 | 0.182 |
| **Females** | n=8 | n=6 |  |
| 16:0Cer | 12.0 ± 2.7 | 11.4 ± 2.2 | 1.000 |
| 18:0Cer | 9.1 ± 1.8 | 8.5 ± 1.8 | 1.000 |
| 20:0Cer | 14.7 ± 3.1 | 14.4 ± 2.8 | 1.000 |
| 22:0Cer | 19.0 ± 3.3 | 20.2 ± 5.0 | 1.000 |
| 24:0Cer | 8.3 ± 1.8 | 8.7 ± 1.5 | 1.000 |
| 24:1Cer | 10.2 ± 2.2 | 10.1 ± 2.1 | 1.000 |
| Total ceramides | 73.3 ± 14.5 | 73.3 ± 14.9 | 0.999 |

Ceramide content, determined by LC/MS, in LV homogenates from 6-month old male and female offspring of control (Con) and obese (Ob) dams. Con and Ob groups compared by Student's t-test with Holm-Sidak multiplicity correction. Mean ± SEM.

## SUPPLEMENTARY METHODS

### Echocardiography

Mice were anaesthetised (2% isoflurane, inhaled), placed in dorsal recumbency on a heated mat maintained at 37°C and hair removed from the thorax using a depilating cream. The heart was imaged in the parasternal short axis, at the level of the papillary muscle, by a trained operator blinded to the treatment group of the animal, using the Vevo 2100 system (VisualSonics). M-mode images of the left ventricle were collected across at least four consecutive cycles and used to measure ventricular wall thicknesses and chamber diameter in systole and diastole. Left ventricular internal volumes at systole and diastole were determined using the leading-edge method then ejection fraction and fractional shortening were calculated as indices of systolic function. Doppler velocimetry was used to determine peak mitral inflow in early (E) and late (A) diastole whilst tissue Doppler was used to determine peak ventricular wall displacement at the level of the mitral annulus, again in early (E’) and late (A’) diastole. E/A, E’/A’ and E/E’ ratios were calculated as indices of ventricular diastolic function. Mice were recovered from anaesthesia following echocardiography and returned to their home cage.

### RNA sequencing

Sequencing libraries with unique barcodes were constructed from 100 ng of total RNA using the KAPA Stranded mRNA-Seq kit (Kapa Biosystems, Wilmington, MA) according to the manufacturer’s protocol. Individual cDNA libraries were quantified by qPCR. Pooled libraries were used to generate clusters by cBot with version 3 reagents (Illumina, San Diego, CA). Multiplex paired-end (2 x 100 base) sequencing was performed on the HiSeq 2500 Sequencing System with version 3 SBS chemistry (Illumina). Sequencing reads were demultiplexed using the CASAVA pipeline (Illumina) and then aligned to the *Mus musculus* reference genome (mm10) using STAR version 2.5.3a in Partek Flow (Partek, St. Louis, MO). Aligned reads were quantified using the Expectation/Maximization algorithm in Partek Flow with RefSeq transcripts from NCBI annotation release 84. Transcripts with zero read counts across all samples were removed prior to performing normalization of read counts using the Trimmed Mean of M-values method ^46^. Afterwards, transcripts with at least 5 read counts across all samples were included for comparing expression in Ob vs. Con, separately in male and female fetuses. The Gene-Specific Analysis in Partek Flow was used for differential expression analysis, in which the best statistical model was identified for each transcript based on the normalized read counts of that transcript and the best model was used to produce the fold change, P values and P values adjusted by the method of Benjamini and Hochberg ^47^.

### Ingenuity Pathway Analysis

Differentially expressed mRNAs were functionally annotated *in silico* using Ingenuity Pathway Analysis (IPA) software (Qiagen). Downstream biological processes predicted to be affected by maternal obesity were identified using an unbiased approach based on significant enrichment with differentially expressed genes. The direction of expression change of mRNAs within each gene set was used to predict overall activation status of the pathway or function, by calculating a z-score, such that negative z-scores represented inhibited pathways and positive z-scores represented activated pathways. Functions were filtered for further investigation based on a threshold of z-score > |1.7| and ranked by significance level (P-value). Downstream functions commonly affected by maternal obesity in both female and male fetuses were identified using unsupervised comparative analysis in IPA. Unsupervised upstream regulator analysis within IPA was also used to identify key molecules (e.g. transcription factors) predicted to cause the observed transcriptional effects of multiple differentially expressed genes.

### Targeted lipidomic analyses

For fatty acid quantification, frozen ventricular tissue was homogenised in hepes-tris buffered saline solution then lipids were extracted using a liquid-liquid extraction method, as described (Chassen et al, 2018). Briefly, an aliquot of ventricle homogenate was deproteinised by addition of methanol, with vortexing, then centrifuged (500g, 15 min, at room temperature). The supernatant solution was transferred to a glass vial, water and dichloromethane were added to extract lipids, vortexed and centrifuged. The polar lipid lower layer was separated and dichloromethane extraction was repeated with the upper aqueous layer. Combined lipid layers were dried under N2, resuspended in ethanol then spiked with an internal FA standard solution. Lipids were saponified with 1M NaOH at 90°C for 1hr, neutralised, extracted into isooctane and derivatized using pentafluorobenzyl bromide and diisopropylethalamine in acetonitrile. The final fatty acid extract was resuspended in isooctane for GC-MS analysis. Samples were separated on a HP-5MS capillary column (30m, 0.25mm, 0.10mm film thickness, Agilent), subjected to mass spectrometry then identified and quantified based on *m/z* ratios and peak heights, respectively ^48^.

For quantification of ceramides and diacylglycerol (DAG) species, heart samples homogenized in 900 µL water and an aliquot (20 µL) taken for protein concentration. Methanol (900 µL) was added to the homogenized sample (750 µL). After the addition of 15:0/18:1(d7)-DAG (80 pmol) and 12:0-ceramide (80 pmol) as internal standards, lipid extraction was performed by the addition of methyl-*tert*-butyl ether (3 mL) according to Matyash et al ^49^. The organic phase was dried under a stream of nitrogen gas and resuspended in 400 µL of a mixture of 70/30 (v/v) hexane:methylene chloride. Samples were injected into an HPLC system connected to a triple quadrupole mass spectrometer (Sciex 2000 QTRAP, Framingham, MA) and normal phase chromatography with a HILIC column (100x2.1 mm, Kinetex HILIC 2.6 µm, Phenomenex) was used to separate lipids by class ^50^. Mass spectrometric analysis was performed in the positive ion mode using multiple-reaction monitoring (MRM) of DAG and ceramide molecular species and the internal standards. Quantitation was performed using stable isotope dilution with a standard curve for DAGs and ceramides and results were normalized to protein content. All lipid measurements were expressed relative to tissue protein content, determined by bicinchoninic acid assay.

